# Population structure and dispersal across small and large spatial scales in a direct developing marine isopod

**DOI:** 10.1101/2020.03.01.971333

**Authors:** William S. Pearman, Sarah J. Wells, Olin K. Silander, Nikki E. Freed, James Dale

**Author notes:** Contributed equally (William S. Pearman, Sarah J. Wells).

## Abstract

Marine organisms generally exhibit one of two developmental modes: biphasic, with distinct adult and larval morphology, and direct development, in which larvae resemble adults. Developmental mode is thought to significantly influence dispersal, with direct developers expected to have much lower dispersal potential. However, in contrast to our relatively good understanding of dispersal and population connectivity for biphasic species, comparatively little is known about direct developers. In this study, we use a panel of 8,020 SNPs to investigate population structure and gene flow for a direct developing species, the New Zealand endemic marine isopod *Isocladus armatus*. On a small spatial scale (20 kms), gene flow between locations is extremely high and suggests an island model of migration. However, over larger spatial scales (600km), populations exhibit a clear pattern of isolation-by-distance. Because our sampling range is intersected by two well-known biogeographic barriers (the East Cape and the Cook Strait), our study provides an opportunity to understand how such barriers influence dispersal in direct developers. Our results indicate that *I. armatus* exhibits significant migration across these barriers, and suggests that ocean currents associated with these locations do not present a barrier to dispersal. Interestingly, we do find evidence of a north-south population genetic break occurring between Māhia and Wellington, two locations where there are no obvious biogeographic barriers between them. We conclude that developmental life history largely predicts dispersal in intertidal marine isopods. However, localised biogeographic processes can disrupt this expectation.

## Introduction

### Life history influences genetic population structure

A species’ developmental life history has a significant effect on population connectivity and dispersal. For marine organisms, developmental life histories are generally classified in one of two ways. Direct developers have juveniles that resemble adults, lacking a pelagic larval phase, and are therefore often hypothesised to have limited dispersal ability relative to their biphasic counterparts. In contrast, biphasic species exhibit a pelagic larval stage have different morphologies from adults and generally disperse large distances via ocean currents (Cowen & Sponaugle, 2009; Puritz et al., 2017; Simpson et al., 2014). Consequently, direct-developing species tend to show greater population genetic structuring than biphasic species (Ayre et al., 2009; McMillan et al., 1992; Pelc et al., 2009; Waples, 1987). However, some species do not conform to this expected pattern (Ayre et al., 2009; Puritz et al., 2017; Shanks, 2009; Winston, 2012).

Dispersal (and thus population structure) is also affected by biogeographic barriers, and the effects of these barriers can themselves depend on the life history of the organism, thus life history and habitat availability jointly act to influence population structure (Ayre et al., 2009). However, habitat availability may be a more important determinant for species with reduced dispersal ability (i.e. direct developers). However, in some cases the lack of nearby suitable habitat appears to limit dispersal in biphasic species more than direct developers (Ayre et al., 2009). Indeed, cases where direct developers exhibit greater dispersal than predicted are not uncommon within intertidal organisms (Ayre et al., 2009; González-Wevar et al., 2018; Wells & Dale, 2018; Yoshino et al., 2018). Therefore, population connectivity cannot be entirely predicted by life history or the knowledge of biogeographic barriers. As a result, it is important to carefully quantify population structure and gene flow in direct developing marine species across a range of spatial scales and biogeographic contexts. This will help elucidate the complex relationships between life history, biogeography, and dispersal.

### Population Genetics in *Isopoda*

Marine isopods offer a tractable model system for investigating the dispersal potential and population structure of direct-developing species. Many isopod species are abundant and easily sampled in intertidal zones across extensive geographic ranges. Nevertheless, the forces acting to maintain population genetic structure in marine isopods are not well understood. Some species exhibit genetic structure over small spatial scales, on the order of tens of kilometres or less (e.g *Idotea chelipes* (Jolly & Rogers, 2003), *Austridotea lacustris* (McGaughran et al., 2006), and *Jaera albifrons* (Piertney & Carvalho, 1994)). This is congruent with the hypothesis of reduced dispersal in direct developers, and may be responsible for the widespread occurrence of multiple cryptic species of isopods (*Ligia* and *Tylos spp.*) on the Southern Californian coastline (Hurtado et al., 2010, 2013; Markow & Pfeiler, 2010).

In contrast, other species of isopods exhibit transoceanic distributions, indicative of historically high dispersal rates. One example is *Sphaeroma terebrans*, which is distributed across both the Atlantic and Indian Oceans. The apparently high dispersal rate and wide distribution of this wood-boring isopod may be result of its reliance on “rafting” for dispersal (a process in which individuals use large rafts of algae or other floating debris to disperse) (Baratti et al., 2011). The contrast between *S. terebrans* and *Ligia* and *Tylos spp*. suggests that life history (i.e. whether species are direct developers or not) may not always be a good predictor of population structure in isopods.

Here, we investigate the New Zealand-wide population genetic structure of the intertidal isopod *Isocladus armatus*. Endemic to New Zealand, this massively colour polymorphic species is found in abundance on semi-sheltered rocky shorelines throughout the country (Jansen, 1971). A previous study concluded that this direct-developing isopod species can exhibit high dispersal rates because no significant population divergence was found between two sites separated by 11km of coastline (Wells & Dale, 2018). In contrast, strong population genetic structure was evident on a much larger scale (1,000kms). Therefore, it is unclear at precisely what spatial scale this population structure begins to break down, and how gene flow is affected by factors such as distance or the presence of biogeographic barriers.

In this study, we use genotyping-by-sequencing (GBS) to resolve population genomic structure over a range of spatial scales. By quantifying population structure in this species, we can identify the influence of geography (both distance and seascape features) on gene flow and dispersal.

## Methods

### Sample Collection

We collected specimens of *Isocladus armatus* in June, 2018, from around the North Island, New Zealand, from locations where *I. armatus* had previously been recorded (Hurley & Jansen, 1977; *INaturalist.org*, n.d.). These sites were at Stanmore Bay, Browns Bay, Opito Bay (Coromandel), Mt Maunganui, Māhia Peninsula, and Wellington (Fig. 1). At each site we collected a minimum of 32 individuals, up to a maximum of 48, and attempted to sample individuals as equally as possible amongst colour morph and sex. Where possible, we collected specimens larger than 5mm in order to ensure enough DNA could be extracted for sequencing (see Supplementary Material for details). The maximum distance between individuals collected at any site did not exceed 30 m. Samples were stored at −80°C in 100% ethanol until extraction. We have included previously collected samples from 2015 (see (Wells & Dale, 2018) for sampling methods) to increase the number of sampled individuals and the range of location. In addition, the temporal structure of this sampling may yield insight into the short-term changes in allele frequencies within a population.

**Figure 1.**
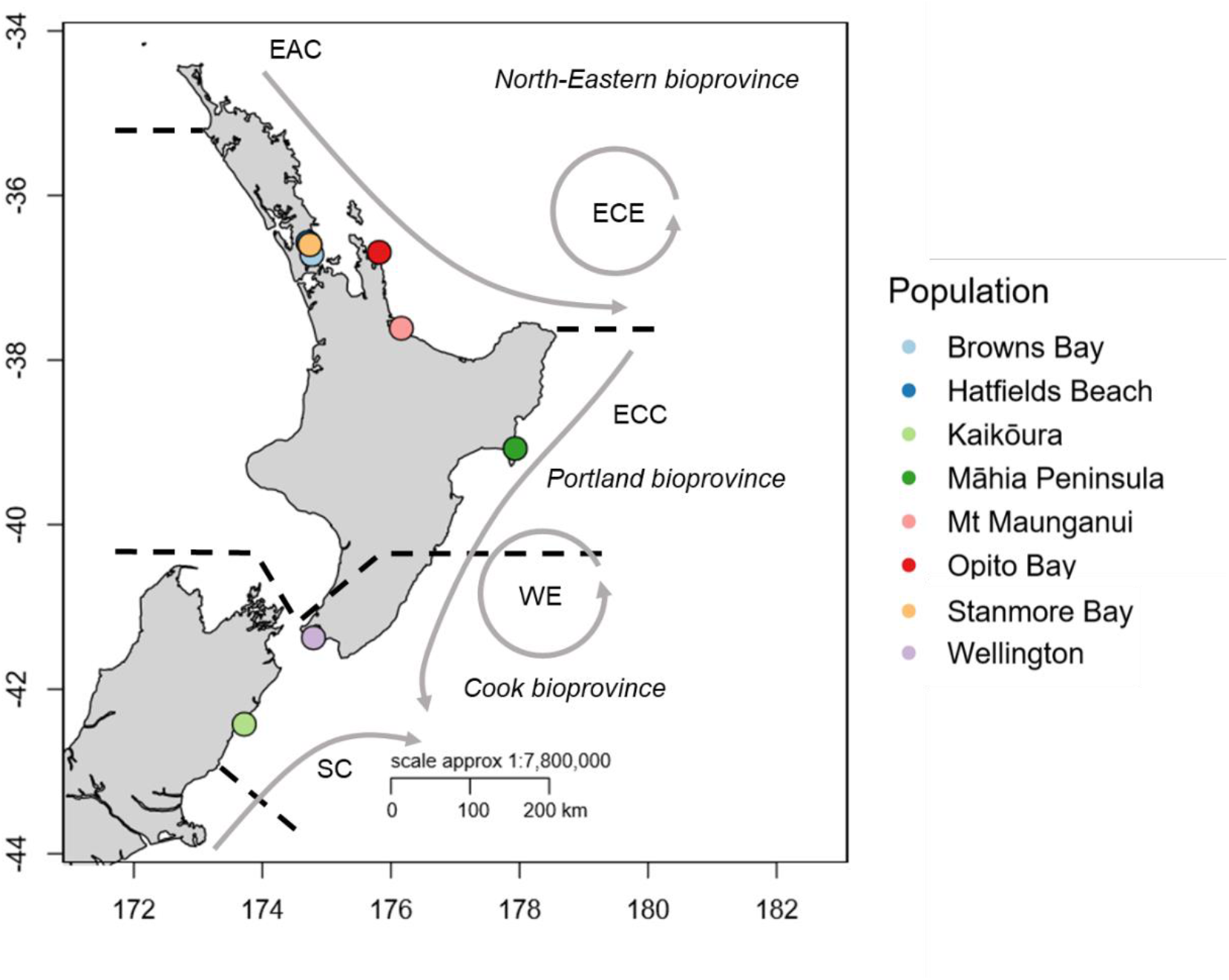
Map of sampling localities within New Zealand (coloured dots). The prevailing ocean currents and biogeographic breaks proposed by Shears et al (2008) are also indicated. Black dashed lines and unabbreviated labels indicate bioprovinces. Grey arrows indicate currents, abbreviated as: EAC (East Auckland Current), ECE (East Cape Eddy), ECC (East Cape Current), WE (Wairarapa Eddy), and SC (Southland Current).

### DNA Extraction

We extracted DNA following a modified Qiagen DNEasy Blood and Tissue protocol from Wells and Dale (2018). Briefly, we used 178 μl of 0.5M EDTA and 22 μl of 20% SDS in each extraction (rather than varying volumes by weight of tissue). Additionally, we eluted DNA from the spin column three times, using 50 μl of nuclease-free water for each. We let the eluent sit on the column for 15 minutes before centrifugation for one minute at 7,000 rcf.

### Data Collection and Processing

DNA samples were processed by Diversity Arrays Technology (DArT) Ltd (Canberra, Australia) using DArTseq, a genotyping-by-sequencing (GBS) approach. The methodology behind this, including restriction enzyme choice are described in detail in Wells and Dale (2018). DArT performs SNP calling using a proprietary pipeline. SNPs are only called if both homozygous and heterozygous genotypes can be identified.

We analysed the dataset provided by diversityArrays together with the data from Wells and Dale (2018). To ensure the datasets were compatible, we filtered each dataset separately based on the conditions described below using the R packages dartR (Gruber et al., 2018) and radiator (Gosselin, 2019). We then used only the loci shared across both datasets for the remainder of the analyses.

We required SNPs to have a call rate ≥ 0.9 (where for at least 90% of individuals a genotype was identified), a minor allele count of at least 3, observed heterozygosity > 0.5, minimum depth of > 5X and maximum depth of 50X (excessively high coverage may suggest duplicated genome elements which could confound further analyses). If we found multiple SNPs on the same read, we removed the SNP with lowest replicability (based on the number of technical replicates that resulted in the same allele being called). Finally, where appropriate for further analysis, we removed SNPs outside of Hardy-Weinberg Equilibrium or that we inferred as being under selection. Specific details of why these filters were implemented and how are outlined in the Supplementary Material. One individual from Browns Bay was excluded from all analyses, as this sample had a very high number of SNPs missing across all loci (93%).

### Data Analysis

We calculated F-statistics using StAMPP (Pembleton et al., 2013). We used F_st_ as the primary measure of genetic differentiation because F_st_ remains relatively robust for biallelic markers such as SNPs (Whitlock, 2011).

We conducted principal component analyses (PCA) using the R package adegenet (Jombart & Ahmed, 2011). In order to understand the correspondence between the principal components and geography, we performed a Procrustes transformation of the first two principal components using MCMCpack in R (Martin et al., 2011). Procrustes transformations scale, stretch, and rotate the PCA in order to minimize the differences between two matrices (in this case, the difference between principal components and geographic coordinates).

We implemented clustering analyses using STRUCTURE 2.3.4 (Falush et al., 2003) performed with all populations (including the repeated samples of Stanmore Bay). For these analyses, an admixture model was assumed. We ran the Markov Chain Monte Carlo simulations with 100,000 iterations and a burn-in of 50,000. We conducted ten replicates of each run, and varied K from 2 through 9. Final population inference was performed by consolidating the results for each level of *K* in CLUMPP (Jakobsson & Rosenberg, 2007). Additionally, we performed a separate STRUCTURE analysis on the Auckland populations with the implementation of the locprior model at a *K* of 3, in order to test for fine-grain population structure within Auckland. Due to concerns regarding the inferences made when defining *K,* we chose to present a range of realistic values for *K* (Lawson et al., 2018; Pritchard et al., 2010; Verity & Nichols, 2016).

We tested for Isolation-by-Distance by conducting a Mantel test using the R package vegan (Oksanen et al., 2010). We used Slatkin’s linearized Fst matrix (transformed using 1/1-F_st_ (Rousset, 1997)) and an overwater distance matrix for this test. We calculated overwater distance using the marmap (Pante & Simon-Bouhet, 2013) and fossil (Vavrek, 2011) R packages, finding the minimum distance between populations around the coast within a depth range of 150 m.

## Results

To examine population structure in *I. armatus* populations, we isolated 261 individuals distributed across 8 populations across New Zealand (**Fig. 1**). We obtained DArTseq SNP data for these 261 individuals, which identified 78,927 SNP loci as being polymorphic. After stringent filtering (see **Methods** and **Supplementary Material**), 8,020 SNPs remained (**Supplementary Table 2**).

We first used this filtered SNP data to test for population structure using F statistics (see **Methods**). We found evidence of genetic structure (likely due to decreased rates of genetic exchange) between populations that were more than 18 km apart (Browns Bay and Stanmore Bay), with F_st_ values ranging from 0.06 to 0.25 (**Table 1**).

**Table 1.**
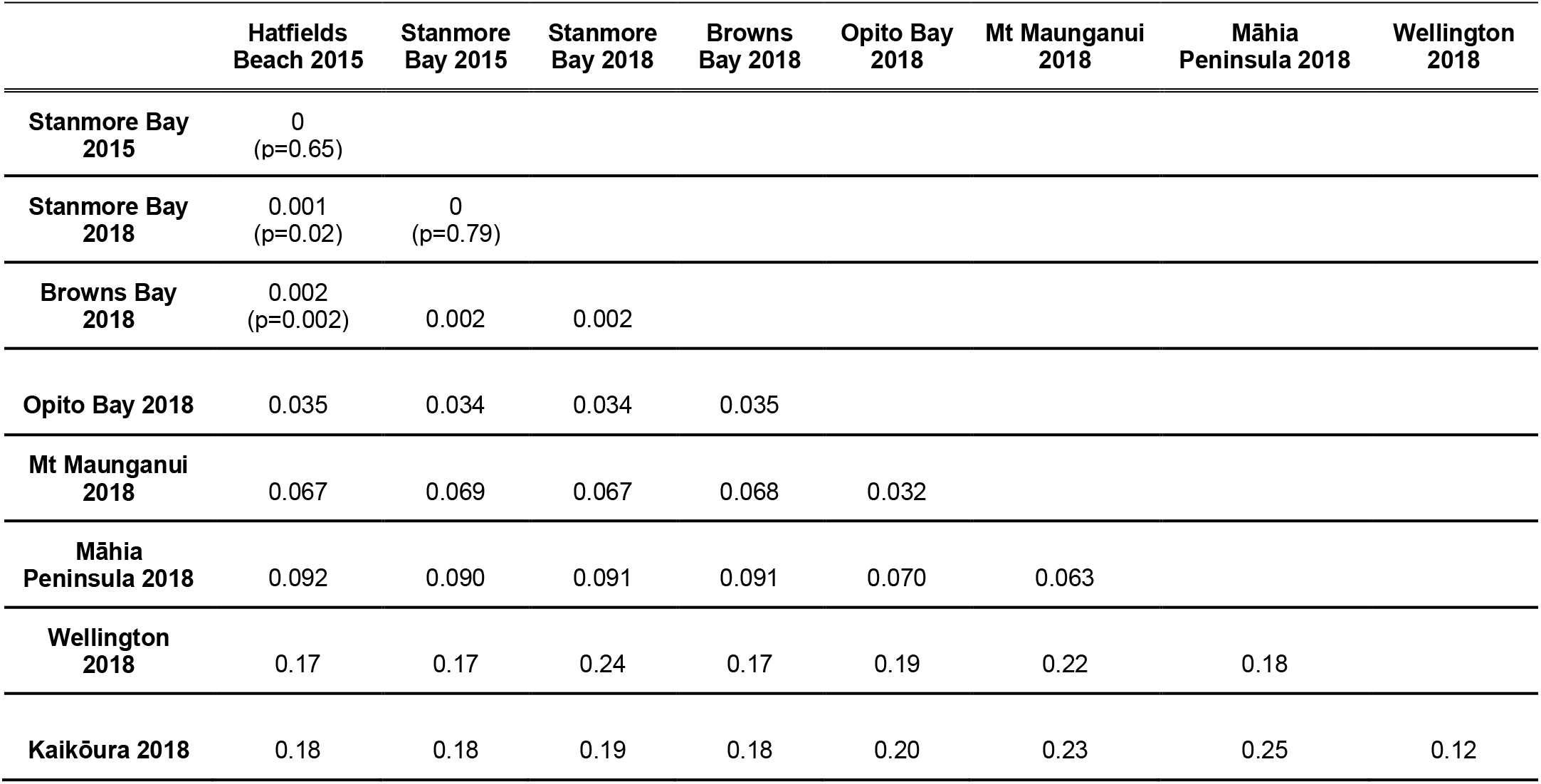
Population F_st_ values. There is evidence of weak population structure within some of the Auckland populations (Hatfields Beach and Stanmore Bay with Browns Bay). However, population structure between all other populations is strong, with a maximal F_st_ of 0.25 between Kaikōura and the Māhia Peninsula. Unless otherwise indicated, all p-values are less than 0.0001.

|  | Hatfields Beach 2015 | Stanmore Bay 2015 | Stanmore Bay 2018 | Browns Bay 2018 | Opito Bay 2018 | Mt Maunganui 2018 | Māhia Peninsula 2018 | Wellington 2018 |
| --- | --- | --- | --- | --- | --- | --- | --- | --- |
| Stanmore Bay 2015 | 0<br>(p=0.65) |  |  |  |  |  |  |  |
| Stanmore Bay 2018 | 0.001<br>(p=0.02) | 0<br>(p=0.79) |  |  |  |  |  |  |
| Browns Bay 2018 | 0.002<br>(p=0.002) | 0.002 | 0.002 |  |  |  |  |  |
| Opito Bay 2018 | 0.035 | 0.034 | 0.034 | 0.035 |  |  |  |  |
| Mt Maunganui 2018 | 0.067 | 0.069 | 0.067 | 0.068 | 0.032 |  |  |  |
| Māhia Peninsula 2018 | 0.092 | 0.090 | 0.091 | 0.091 | 0.070 | 0.063 |  |  |
| Wellington 2018 | 0.17 | 0.17 | 0.24 | 0.17 | 0.19 | 0.22 | 0.18 |  |
| Kaikōura 2018 | 0.18 | 0.18 | 0.19 | 0.18 | 0.20 | 0.23 | 0.25 | 0.12 |

We next quantified population structure using principal component analysis, which identifies the combinations of SNP loci that vary the most between individuals. We found that PC1 accounted for 19.5%, PC2 for 3.65%, and PC3 for 2.11% of the variance in SNP allele frequency (**Figs. 2A** and **2B**). PC1 primarily delineated the southern Wellington and Kaikōura populations from the other populations. PC2 primarily differentiated between the northern populations, and revealed three potential cases of migration between Mt Maunganui and the Māhia Peninsula (**Fig. 2A**). Finally, PC3 differentiated the southern populations, Wellington and Kaikoura. All Auckland populations clustered together, suggesting these individuals form one large population. We found no difference between the temporal samples from Stanmore Bay, indicating that allele frequency variation over short timescales is relatively stable.

**Figure 2.**
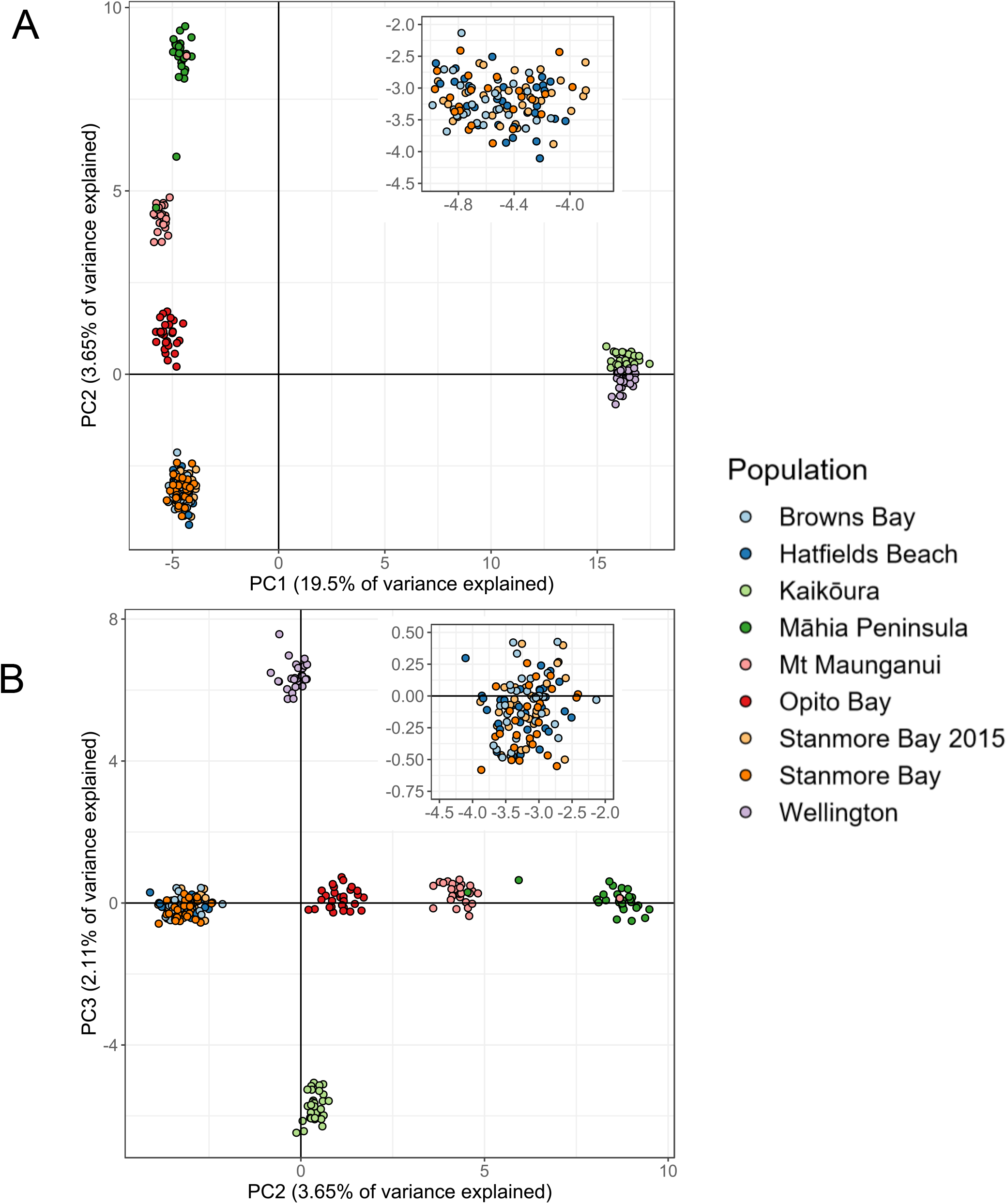
Principal Component Analysis (PCA) indicates strong location-dependent population structure. PC1 (19.5% of the variance) largely differentiates the individuals in southern populations from those in the northern populations (i.e. Wellington and Kaikōura from the rest). PC2 (3.65% of the variance) differentiates between the northern populations. Finally, PC3 differentiates the populations within the southern group. Insets show the Auckland populations, and indicate minimal evidence for structure within them. Three potential recent migrant individuals are apparent, two from the Māhia population (dark green points) to the Mt Maunganui population (pink points), and one from the Mt Maunganui population to the Māhia population, visible in panel (B) within the dark green points.

A Procrustes transformation of the first two principal components of the PCA onto geographical space revealed that the spacing of each genetic cluster was strikingly concordant with the locations in geographical space, with the exception of Māhia and Wellington (**Fig. 3**).

**Figure 1.**
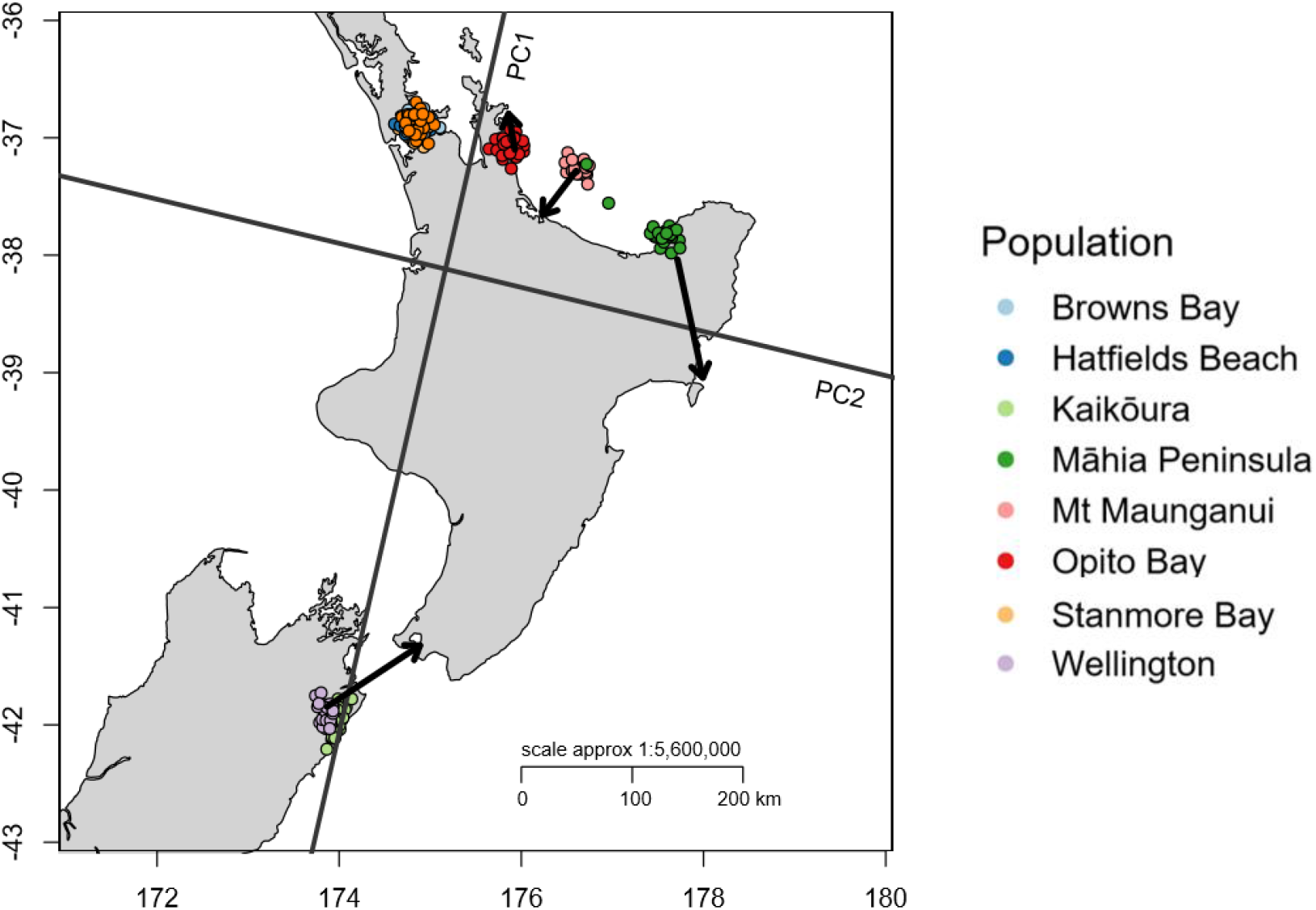
Procrustes transformation of PC1 vs PC2 onto geographical coordinates. The transformation indicates a strong correspondence between sampling location and genotype distance in principal component space. The arrows point to the location where a population is actually found, while clusters indicate their location in Procrustes-transformed space. The bottom arrow indicates the location of the Wellington population (purple) only.

As a complementary approach to the PCA in quantifying population structure, we implemented a STRUCTURE analysis. This method implements a model-based clustering approach, which probabilistically assigns individuals to one or more populations under an admixture model. The admixture model assigns different proportions of an individual's genome belonging to different ancestral populations (Pritchard et al., 2010). The STRUCTURE analyses showed similar results to the PCA, with all individuals from the Auckland locations consistently clustering together, while Kaikōura and Wellington also always clustered together across all ranges of K (**Fig. 4**). Again we observed no fine-grain structure within Auckland when sampling location was incorporated into the analysis (through use of the locprior model) (**Suppl. Fig. 3**).

**Figure 4.**
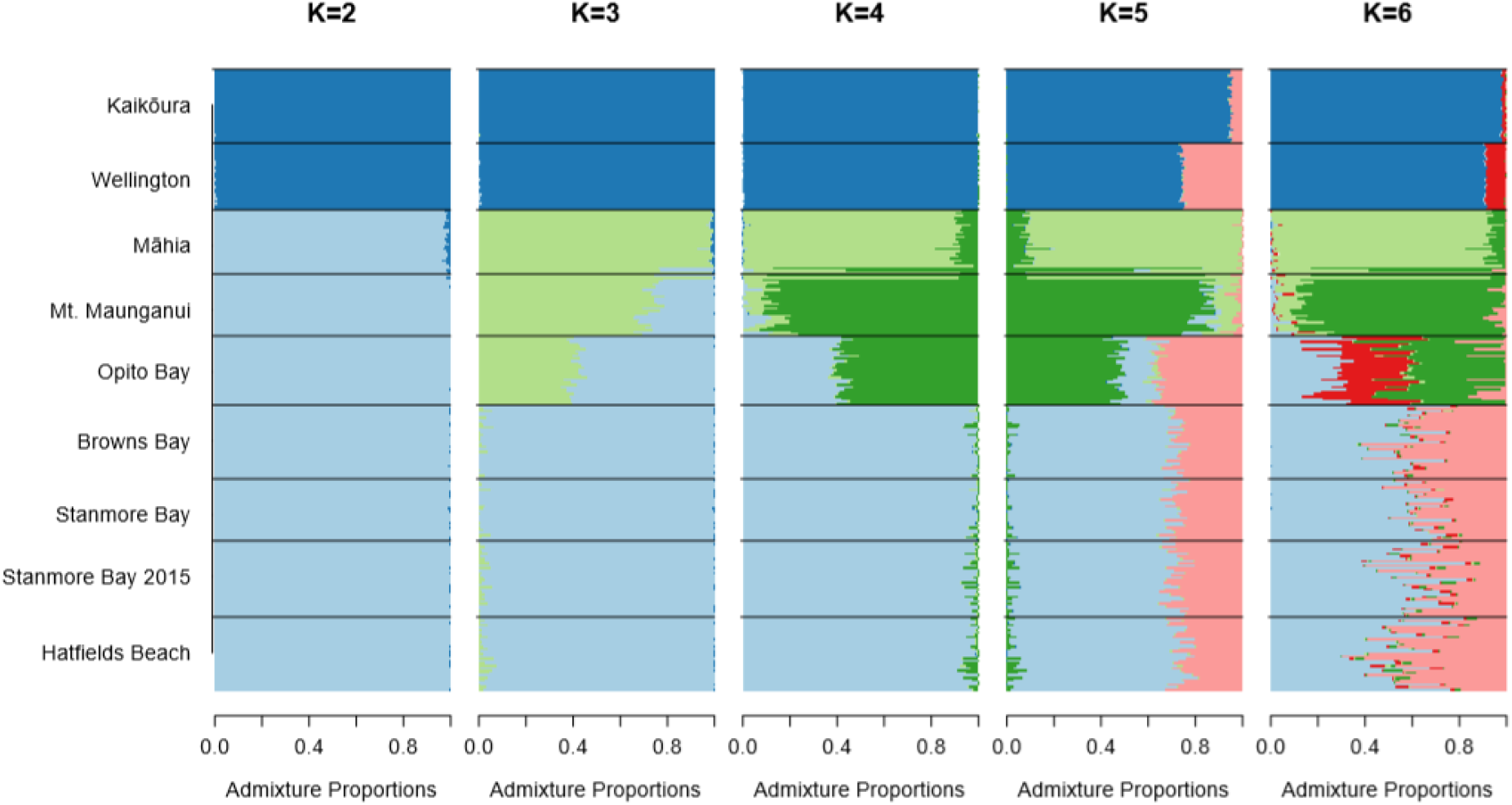
Admixture plots based on Bayesian clustering analyses generated in STRUCTURE. Each horizontal coloured line represents an individual from the locality sampled (indicated on the left). Horizontal black lines designate the junction between population samples. Results are based on 10 replicate runs for each K. At K=4 there are two individuals from Māhia with Mt. Maunganui-like admixture profiles (bottom two bars in the Māhia samples), and one individual from Mt. Maunganui with a Māhia-like admixture profile (top bar in the Mt. Maunganui samples).

As we increased K, additional population genetic structure became apparent. When K=3, Māhia individuals formed a distinct cluster exhibiting high admixture with both Mt. Maunganui and Opito Bay. Increasing K further suggested genetic admixture between all adjacent populations with the exception of Wellington and Māhia. With K=4, there was clear evidence of two Māhia individuals with admixture profiles more similar to Mt. Maunganui individuals, and one Mt. Maunganui individual with a profile similar to Māhia individuals (**Fig. 4**, bottom two individuals in the Māhia sample and top individual in the Maunganui sample, respectively). These correspond to the potential migrants identified in the PCA above. Unexpectedly, with K=5 and K=6, there was evidence of admixture between the northernmost populations (Auckland and Opito Bay) and the southernmost (Wellington and Kaikōura). However, this may be an artefact of overfitting (Evanno et al., 2005).

For all the analyses above (F_st_, PCA, and STRUCTURE), there was clear evidence of population structure. In addition, the Procrustes transformation showed high correspondence between PCA distance and geographic distance (i.e. isolation-by-distance, IBD). In order to further confirm this, we used a Mantel test. This test showed a significant positive correlation between Slatkin’s linearized F_st_ (used as a measure of population structure) and geographic overwater distance (r = 0.83, P = 0.001; **Fig. 5**), again supporting the hypothesis of IBD. This strong positive relationship remained even after excluding the southernmost populations (Kaikōura and Wellington) (r = 0.92, P = 0.001). A molecular analysis of variance also confirmed the existence of genetic structure between the southern populations (Wellington and Kaikōura) and the northern populations, with the majority of between-population variance being explained by this split (see **Supplementary Material**).

**Figure 5.**
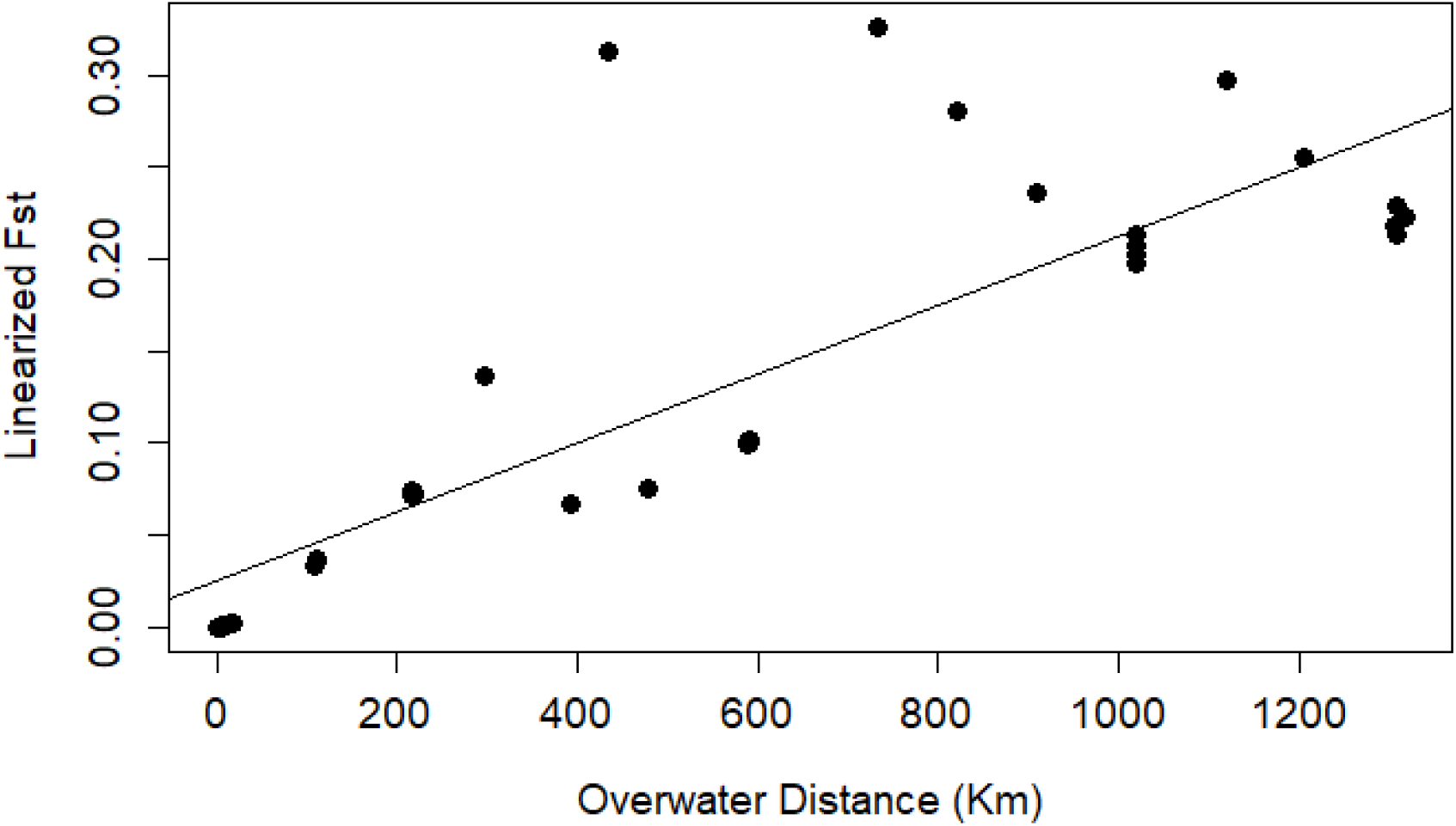
The positive correlation between overwater distance and Slatkin’s linearized F_st_ suggests a pattern of population genetic structure due to isolation-by-distance. The black line indicates the slope of relationship for just the contrasts between the northern populations (points in the lower left quadrant).

## Discussion

Here we have quantified population genetic structure in a direct-developing marine invertebrate across a wide range of spatial scales. By employing a GBS approach we aimed to increase our power to detect even low levels of population structure.

We found minimal evidence of population structure of *Isocladus armatus* within the Auckland region. Only the F_st_ analysis suggested that there is any population differentiation, between Browns Bay and the other Auckland populations. These sites are separated by approximately 20km, which may be the smallest distance at which population genetic structure starts to occur in this species. Further resolution of this would require more fine-scaled spatial sampling in this region. These results, indicating low levels of population structure, are consistent with previous work in this species (Wells & Dale, 2018), as well as some biphasically-developing marine invertebrates in the Hauraki Gulf surrounding Auckland, such as the native New Zealand sea urchin *Evechinus chloroticus* (Nagel et al., 2015) and the invasive tunicate *Styela clava* (Goldstien et al., 2010).

Over larger spatial scales, *I. armatus* exhibited strong patterns of isolation by distance (IBD). The Procrustes-transformed PCA indicated that the first two principal components were highly concordant with the geographic arrangement of populations. To test the hypothesis of IBD more explicitly, we used a Mantel test, which indicated a strong and almost linear correlation between geographic distance and genetic distance, indicative of a stepping stone model of distribution. To try and estimate specific migration rates, we also performed a BayesAss3 analysis (Mussman et al., 2019). However, the parameter estimates from this analysis were internally inconsistent and, in some cases, had unbounded confidence intervals (see **Supplementary Material**). This suggested that additional sampling would be required to accurately estimate these parameters.

We found evidence for unusually high admixture between Mt. Maunganui and Māhia Peninsula, despite their separation by a well-known biogeographic barrier, the East Cape. The strongest evidence for this is the presence of three individuals that appeared to be recent migrants between these populations. In one of these an individual from Māhia population clustered with the Mt Maunganui in the PCA, while the opposite was the case for the second individual. The third individual was found located between the associated PC clusters. These three individuals also clustered with the other population, or were highly admixed, in the STRUCTURE analysis, supporting the migrant hypothesis. The individual with an intermediate genotype, as indicated by both the PCA and STRUCTURE analyses, argues against these results being due to sample contamination.

The biogeographic break at the East Cape has been shown to affect population structure in a range of species. This includes direct developers such as the anemone *Actinia tenebrosa* and two species of amphipods (Stevens & Hogg, 2004; Veale & Lavery, 2012). Even population structure in biphasic species with larval stages, such as the paua *Haliotis iris* (Will et al., 2011), and the marine gastropod *Buccinulum vittatum* (Gemmell et al., 2018), are affected by the East Cape. However, in contrast to these other direct developers, this break does not appear to strongly affect *Isocladus armatus*, as F_st_ between Mt. Maunganui and Māhia was no greater than F_st_ between Mt. Maunganui and the Auckland area. In addition, the PCA indicated that the Mt. Maunganui population was almost exactly intermediate in genetic similarity between the Māhia and Opito Bay populations, rather than being more closely associated with the Opito Bay population, given their greater proximity. Finally, the STRUCTURE analyses also showed admixture between Māhia and Mt. Maunganui across all values of K.

Instead of the expected north-south genetic break at the East Cape, we found a strong north-south break between Māhia and Wellington. This north-south break is congruent with the placement of a proposed biogeographic region border in this area (Shears et al., 2008) (**Fig. 1**). However, Shears et al. (2008) also proposed a biogeographic region border at East Cape, which is inconsistent with our study. Indeed, our study contrasts with two previously proposed hypotheses of strong genetic breaks associated with East Cape and the Cook Strait. Instead, we observe high gene flow over the East Cape, and genetic clustering of Wellington and Kaikōura populations.

One explanation for the strong genetic break may be the presence of cryptic species within *I. armatus*, which would drive gene flow to near-zero levels. Within New Zealand, strong north-south divergence between populations of the brooding brittle star, *Amphipolis squamata*, has been suggested to be the result of cryptic speciation (Sponer & Roy, 2002). Cryptic species have been frequently observed in isopods (Hurtado et al., 2016; Leese et al., 2008; Markow & Pfeiler, 2010), and the degree of genetic divergence between the northern and southern group in *I. armatus* is similar to that found between other cryptic species of isopods based on mitochondrial DNA (Leese et al., 2008). The potential for cryptic species is further supported by the observation of an individual from Browns Bay (which was excluded from all analyses) that, despite appearing morphologically similar to *I. armatus*, lacked 93% of SNPs that were present in other samples. While missing data is a common feature of reduced representation datasets, excessively high missingness in GBS data has been associated with divisions between species rather than populations (Tripp et al., 2017).

## Conclusion

*Isocladus armatus* exhibits a surprisingly high amount of gene flow across small spatial scales. However at distances greater than 20 km the level of population structure is consistent with the expectation of reduced dispersal in direct developing species, and the presence of IBD. Interestingly, the strongest genetic break we observed was between the Māhia Peninsula and Wellington, with populations forming a clear northern and southern grouping either side of this break. This was unexpected, as other well-known biogeographic barriers – the East Cape and the Cook Strait – appeared to have little effect on population genetic structure. Additional fine-scale sampling across this genetic break would help determine whether differentiation across this break is a result of the existence of cryptic species being present, or the result of a geophysical barrier that prevents dispersal in this region.

## Supplementary Material

### Supplementary Methods

#### Sample Collection

*Isocladus armatus* exhibits a wide range of colour morphs, which we refer to as striped, variegated, white, unpatterned, or other (Wells and Dale 2018). During the collection of individuals to quantify population structure, we collected at least 32 individuals from each location (**Supplementary Table 1**). For cases in which more than 32 samples were collected, we selected samples for DNA extraction and subsequent sequencing based on the size and degree of colour morphotype (less ambiguous morphotypes were chosen preferentially over the more ambiguous ones, **Supplementary Table 1**).

**Supplementary Table 1.** Number of samples from each location grouped by sex and colour morphotype. All sites are from 2018 except where explicitly stated.

| Site | Sex | Striped | Unpatterned | Variegated | White | Other | Sum |
| --- | --- | --- | --- | --- | --- | --- | --- |
| Hatfields Beach 2015 | Female | 3 | 6 | 4 | 2 | 1 | 16 |
| Hatfields 2015 | Male | 0 | 4 | 4 | 4 | 3 | 15 |
| Stanmore Bay 2015 | Female | 4 | 4 | 4 | 4 | 0 | 16 |
| Stanmore Bay 2015 | Male | 4 | 4 | 4 | 4 | 0 | 16 |
| Stanmore Bay | Female | 4 | 4 | 4 | 4 | 0 | 16 |
| Stanmore Bay | Male | 4 | 4 | 4 | 4 | 0 | 16 |
| Browns Bay | Female | 4 | 4 | 4 | 4 | 0 | 16 |
| Browns Bay | Male | 4 | 4 | 4 | 4 | 0 | 16 |
| Māhia Peninsula | Female | 0 | 6 | 4 | 3 | 0 | 13 |
| Māhia Peninsula | Male | 2 | 9 | 5 | 0 | 0 | 16 |
| Mt Maunganui | Female | 1 | 3 | 3 | 4 | 0 | 11 |
| Mt Maunganui | Male | 4 | 4 | 4 | 3 | 0 | 15 |
| Opito Bay | Female | 4 | 6 | 1 | 2 | 0 | 13 |
| Opito Bay | Male | 3 | 6 | 4 | 3 | 0 | 16 |
| Wellington | Female | 0 | 5 | 5 | 5 | 0 | 15 |
| Wellington | Male | 0 | 5 | 5 | 5 | 0 | 15 |
| Kaikōura 2015 | Female | 1 | 7 | 4 | 3 | 0 | 15 |
| Kaikōura 2015 | Male | 1 | 5 | 4 | 0 | 5 | 15 |

#### SNP Filtering

A SNP call rate of ≥ 0.9 was implemented. This ensures that for each locus, a SNP genotype was identified in at least 90% of all individuals. Mean read depths of >5X and <50X were also implemented. While high read depths are often desired in other scenarios, excessively high coverage may suggest duplicated genome elements which could confound analyses. On the other hand, low read depths (<5X) may reduce confidence in the accuracy of an allele identification.

In cases where multiple SNPs were located on the same sequence read, we retained the SNP that had the highest reproducibility (based on the proportion of technical replicates that resulted in the same allele being called). In the event these were equal, then the SNP with the highest amount variation across individuals was retained. Removing these SNPs ensures that polymorphisms known to be tightly linked are removed, while removing the SNP with the lowest variation ensures that the most informative loci are retained.

Finally, we removed loci that we inferred as being under strong selection. Non-neutral loci are expected to exhibit different allele frequency spectra to neutral loci. Loci under strong selection may bias inference of population demographics (Luikart et al., 2003). Loci under selection were identified using BayeScan (Foll & Gaggiotti, 2008), with population as the grouping factor. Loci exhibiting a q-value of ≤ 0.05 were excluded from any further analyses.

While it is common practice to filter loci based on being in Hardy-Weinberg equilibrium (HWE) (Morin et al., 2009; Van Wyngaarden et al., 2017; Waples, 2015; Wells & Dale, 2018), this is not always the best practice. For example, STRUCTURE clusters individuals into populations in Hardy-Weinberg equilibrium (Pritchard et al., 2010). By implementing a filter on HWE, sites informative for population structure would be removed. The only analyses for which a HWE filter was implemented were the F_st_ and the associated mantel test. Loci out of HWE were first identified using the R package *pegas* with Bonferroni correction for multiple testing, and removed using the dartR package.

### Supplemental Analyses

#### Analysis of Molecular Variance

We conducted a hierarchical Analysis of Molecular Variance (AMOVA) using the R packages *ade4* (Dray et al., 2007) and *poppr* (Kamvar et al., 2014). This approach partitions the variance in genetic data among and within each level of a predefined hierarchy (or example, individuals, populations, or large geographically-defined groups) (Excoffier et al., 1992). We tested the significance of the AMOVA by performing 1000 random permutations using the R package *pegas*. We used a hierarchy (in ascending order) of sample, population, and then by the observed north-south division found in our other analyses.

#### Migration Rates

We estimated migration rates between populations using BayesAss3-SNPs (Mussmann et al., 2019; Wilson & Rannala, 2003). We used the BA3-Autotune software developed alongside BayesAss3-SNPs to identify optimal parameters for the analysis (Mussmann et al., 2019). We excluded the 2015 Stanmore Bay population for this analysis to avoid duplicating a population for migration estimates.

We conducted this analysis using 1000 randomly sampled loci, and used the following parameters from BA3-Autotune: allele-frequency mixing of 0.2125, migration mixing of 0.075, and inbreeding mixing of 0.075. We ran the analysis for 10 million iterations, with a burn-in of 5 million iterations.

### Supplementary Results

**Supplementary Table 2.**
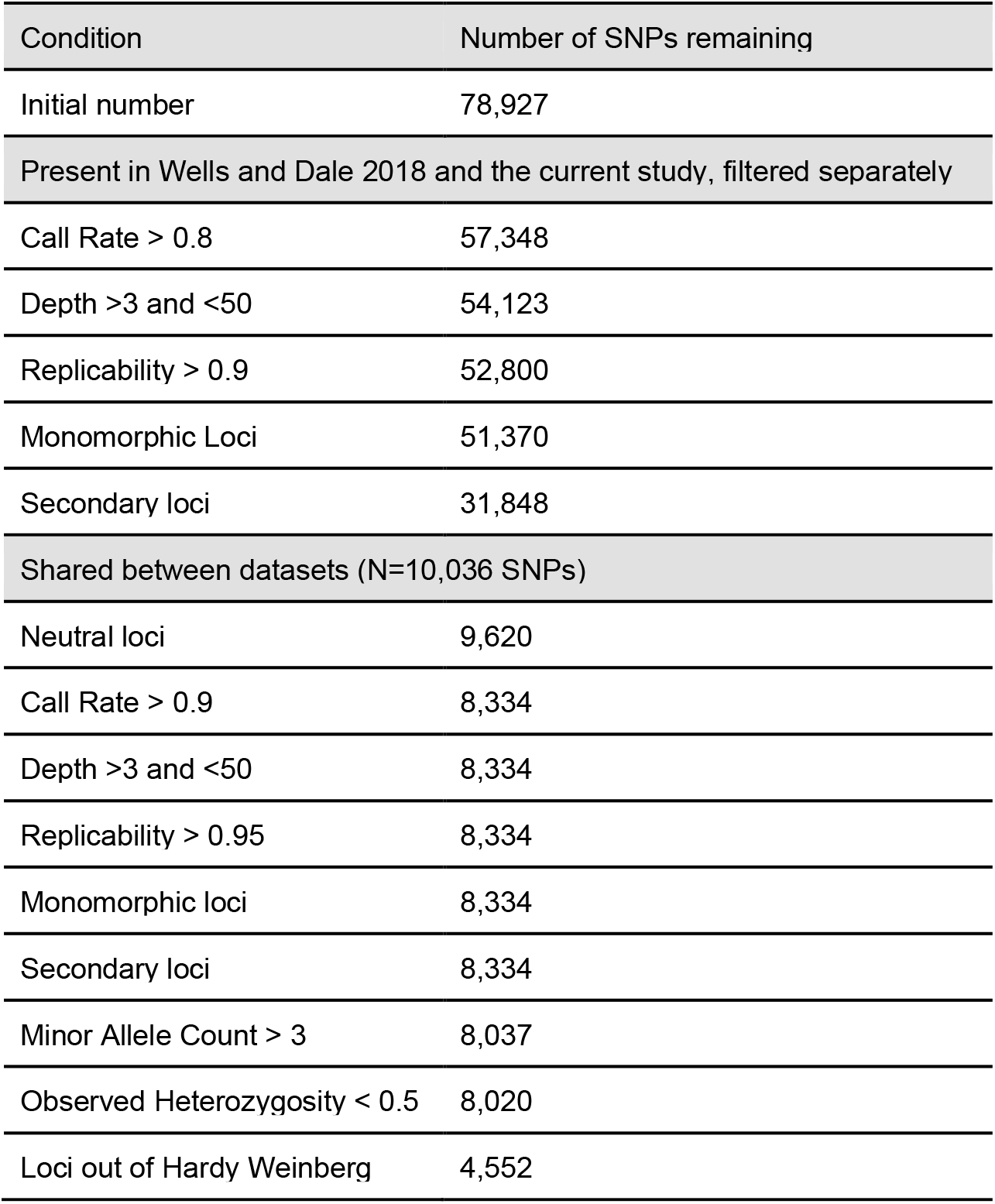
SNP filtering results and the amount of SNPs remaining after each filter. Datasets were first filtered separately and shared SNPs between the datasets were retained for the combined dataset. This was performed to minimise batch effects as a result of samples being sequenced at two different times. Additionally, the increased data output in the 2018 dataset (as a result of the inclusion of more individuals and populations) meant that almost twice as many SNPs were identified in this dataset, than in the 2015 dataset. By retaining shared SNPs the effect of this is minimized. Call rate refers to the proportion of individuals for which a genotype can be identified for the loci.

**Supplementary Figure 1.**
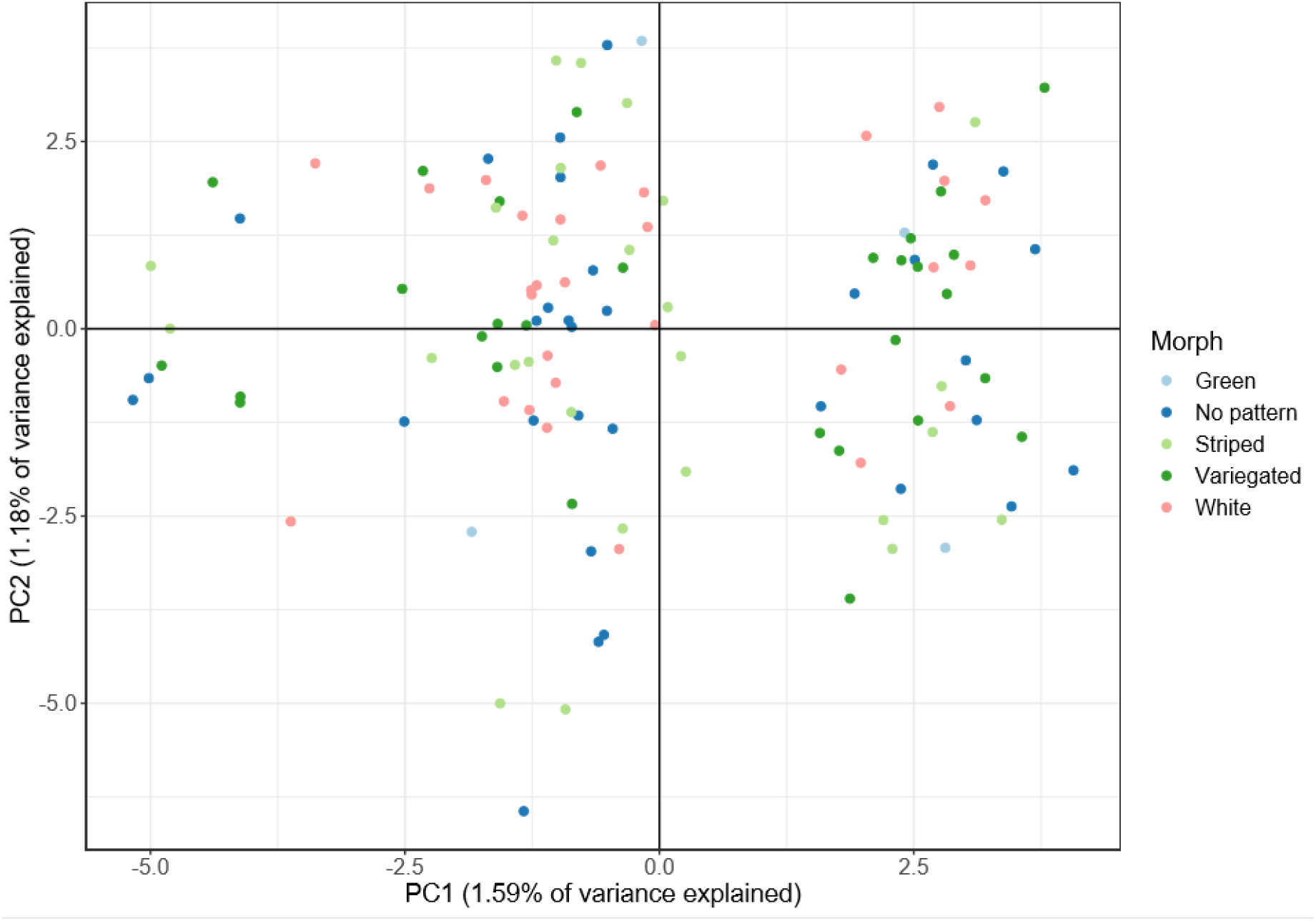
Principal Component Analysis of Auckland populations, coloured by morphotype. No substructuring or stratifying effect of colouration is observed.

**Supplementary Figure 2.**
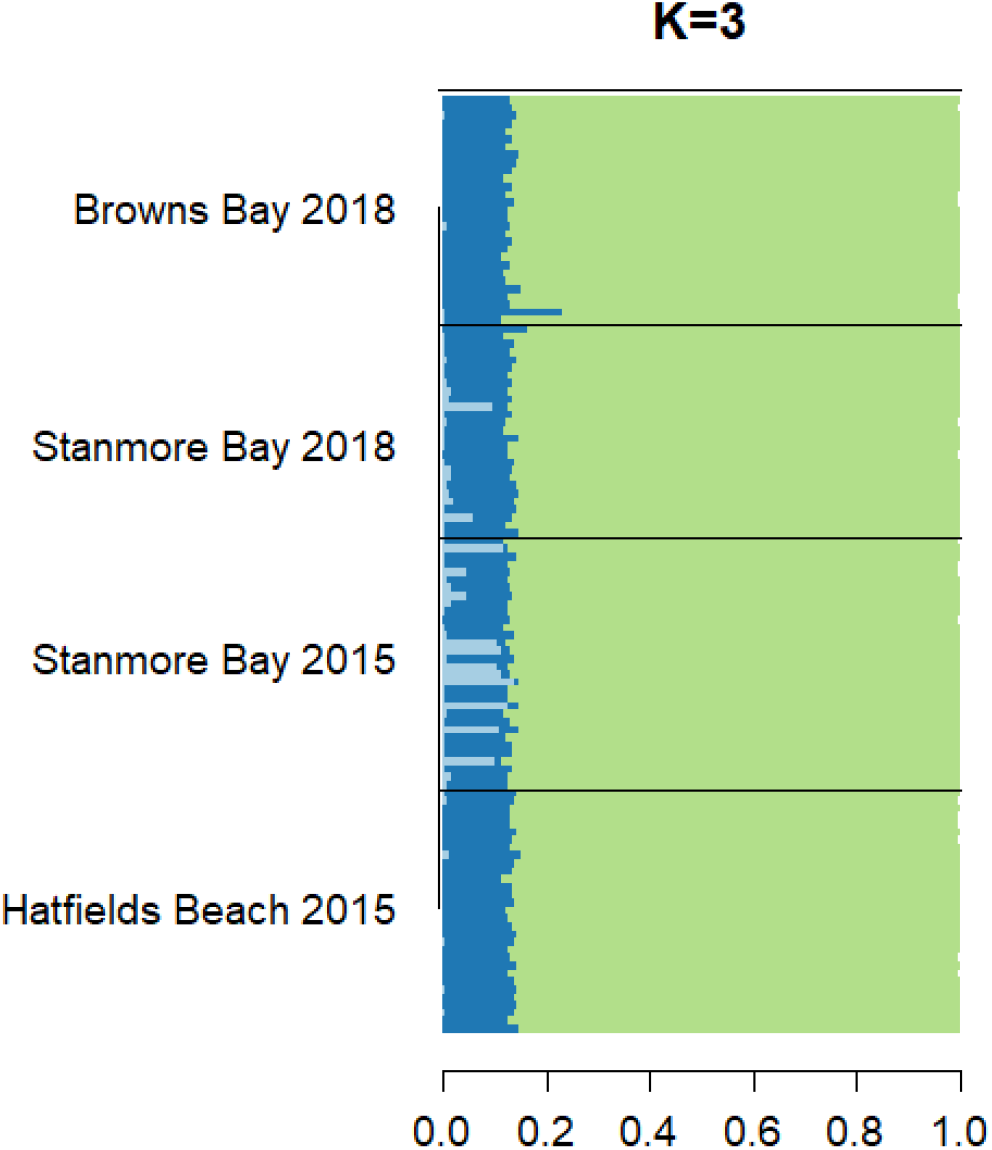
Admixture barplots generated through STRUCTURE and CLUMPP for a value of 3 for K. This analysis was done with the locprior model on just Auckland populations, and was used to identify fine grain structure.

In order to understand the levels at which genetic variation could be attributed, we performed an AMOVA. This analysis indicated some degree of population structure (**Supplementary Table 3**). The majority of genetic variation was within samples (56.2%), followed by regions (28.9%). Between population within region variance was just below 5%.

**Supplementary Table 3.**
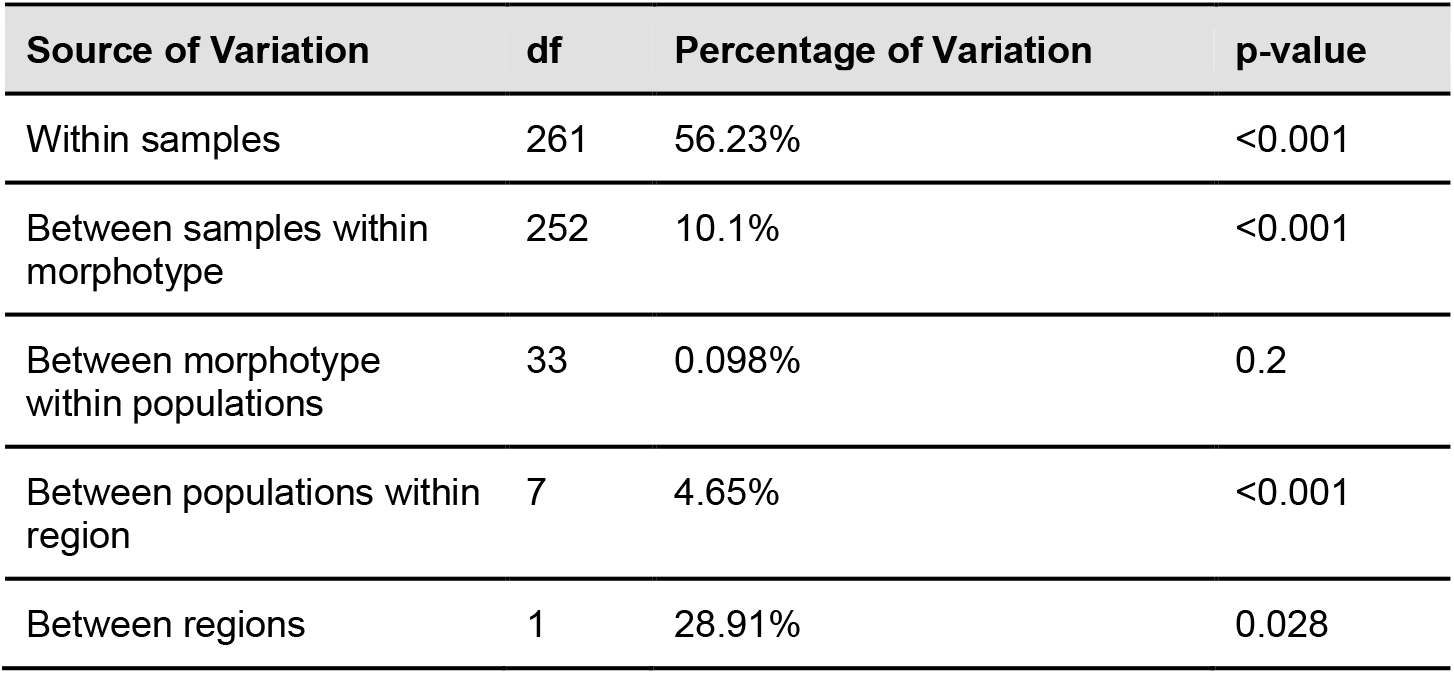
AMOVA analysis shows population structure between populations and regions based on genetic variation from 8,020 SNPs. We defined regions as the north group (all populations excluding Kaikōura and Wellington) and south group (Kaikōura and Wellington).

| Source of Variation | df | Percentage of Variation | p-value |
| --- | --- | --- | --- |
| Within samples | 261 | 56.23% | <0.001 |
| Between samples within morphotype | 252 | 10.1% | <0.001 |
| Between morphotype within populations | 33 | 0.098% | 0.2 |
| Between populations within region | 7 | 4.65% | <0.001 |
| Between regions | 1 | 28.91% | 0.028 |

#### Estimation of Migration Rates

We tested for migration events using BayesAss3. This analysis suggested low migration between most populations. These three groups correspond to 1) the Auckland based populations, 2) the East Coast of North Island populations and, 3) the Southern populations, similar to what was found in other analyses. These estimates indicated no migration in or out of the Southern population, but some negligible migration into Wellington from Kaikōura (Fig 6 and Table 3).

**Supplementary Table 4.** BayesAss migration estimates (as proportion of migrant individuals in population). Values in bold are those used in the final analyses and used to create figure 6. Migration rates represent migration from the population on the top into the population on the left, i.e. the upper triangle represents northward migration, while the bottom triangle represents southward migration. Values in brackets represent standard deviations, as estimated by BayesAss. Heatmap colours are based on log10 transformed values.

|  | Hatfields Beach | Stanmore Bay | Browns Bay | Opito Bay | Mt Maunganui | Māhia Peninsula | Wellington | Kaikōura |
| --- | --- | --- | --- | --- | --- | --- | --- | --- |
| Hatfields Beach | 0.675<br>(0.008) | 0.008 (0.008) | 0.267<br>(0.021) | 0.008<br>(0.008) | 0.008 (0.008) | 0.008<br>(0.008) | 0.008 (0.008) | 0.008<br>(0.008) |
| Stanmore Bay | 0.009<br>(0.009) | 0.676 (0.009) | 0.258<br>(0.024) | 0.009<br>(0.009) | 0.009 (0.009) | 0.01<br>(0.009) | 0.01 (0.009) | 0.009<br>(0.009) |
| Browns Bay | 0.008<br>(0.008) | 0.008 (0.008) | 0.917<br>(0.023) | 0.025<br>(0.014) | 0.008 (0.008) | 0.008<br>(0.008) | 0.008 (0.008) | 0.008<br>(0.008) |
| Opito Bay | 0.009<br>(0.008) | 0.009 (0.009) | 0.009<br>(0.009) | 0.925<br>(0.023) | 0.014 (0.011) | 0.009<br>(0.009) | 0.009 (0.009) | 0.009<br>(0.009) |
| Mt Maunganui | 0.01 (0.009) | 0.01 (0.009) | 0.01<br>(0.009) | 0.09<br>(0.027) | 0.833 (0.029) | 0.019<br>(0.013) | 0.009 (0.009) | 0.01 (0.009) |
| Māhia Peninsula | 0.009<br>(0.009) | 0.009 (0.009) | 0.009<br>(0.009) | 0.01<br>(0.009) | 0.037 (0.017) | 0.898<br>(0.025) | 0.009 (0.009) | 0.009<br>(0.009) |
| Wellington | 0.009<br>(0.009) | 0.009 (0.009) | 0.009<br>(0.009) | 0.009<br>(0.009) | 0.009 (0.009) | 0.009<br>(0.009) | 0.926 (0.023) | 0.011 (0.01) |
| Kaikōura | 0.008<br>(0.008) | 0.008 (0.008) | 0.008<br>(0.008) | 0.008<br>(0.008) | 0.008 (0.008) | 0.009<br>(0.008) | 0.008 (0.008) | 0.934<br>(0.021) |

**Supplementary Figure 6.**
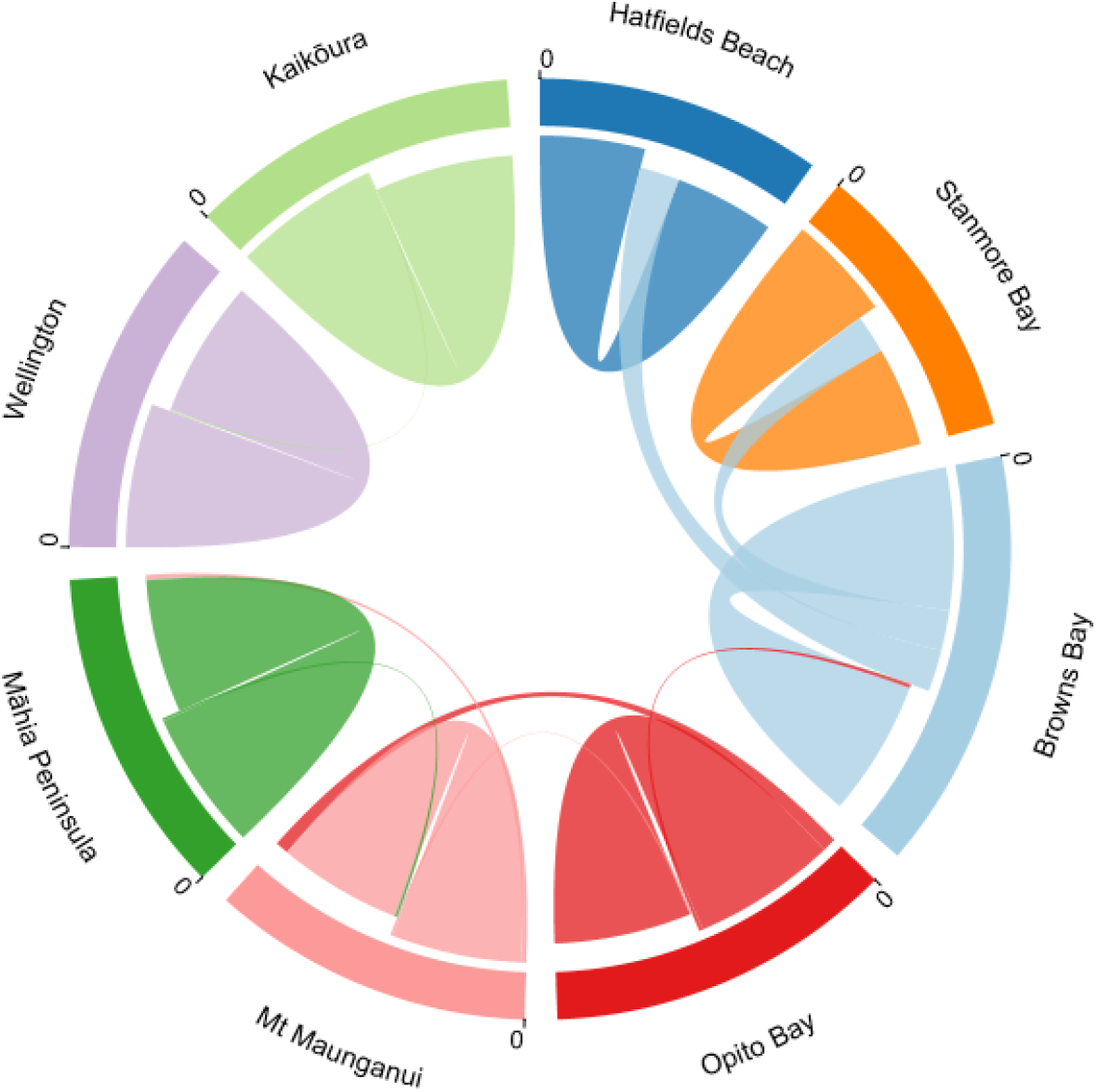
Chord diagram of migration estimates between each population. These estimates represent the proportion of individuals in a population that are estimated to be migrants. The largest arc for each population indicates within population movement, such as a self-recruitment, while the smaller arcs are indicative of between population movement or migration. As a result, wider arcs indicate higher migration rates. Displayed migration rates are those for which the migration rate minus the standard deviation was greater than 0.005.

High migration within the Auckland populations was observed, with estimates of migration rates approximately 0.25, this value represents the proportion of individuals within a population that are estimated to be migrants from the source population. Further south, among the populations along the East Coast of the North Island, migration estimates were consistently higher among adjacent populations than between non-neighbouring populations, suggestive of a stepping stone model of migration (Kimura & Weiss, 1964). These migration rates were relatively low - generally between 0.01 and 0.02 (Supplementary Table 3).

**Supplementary Figure 7.**
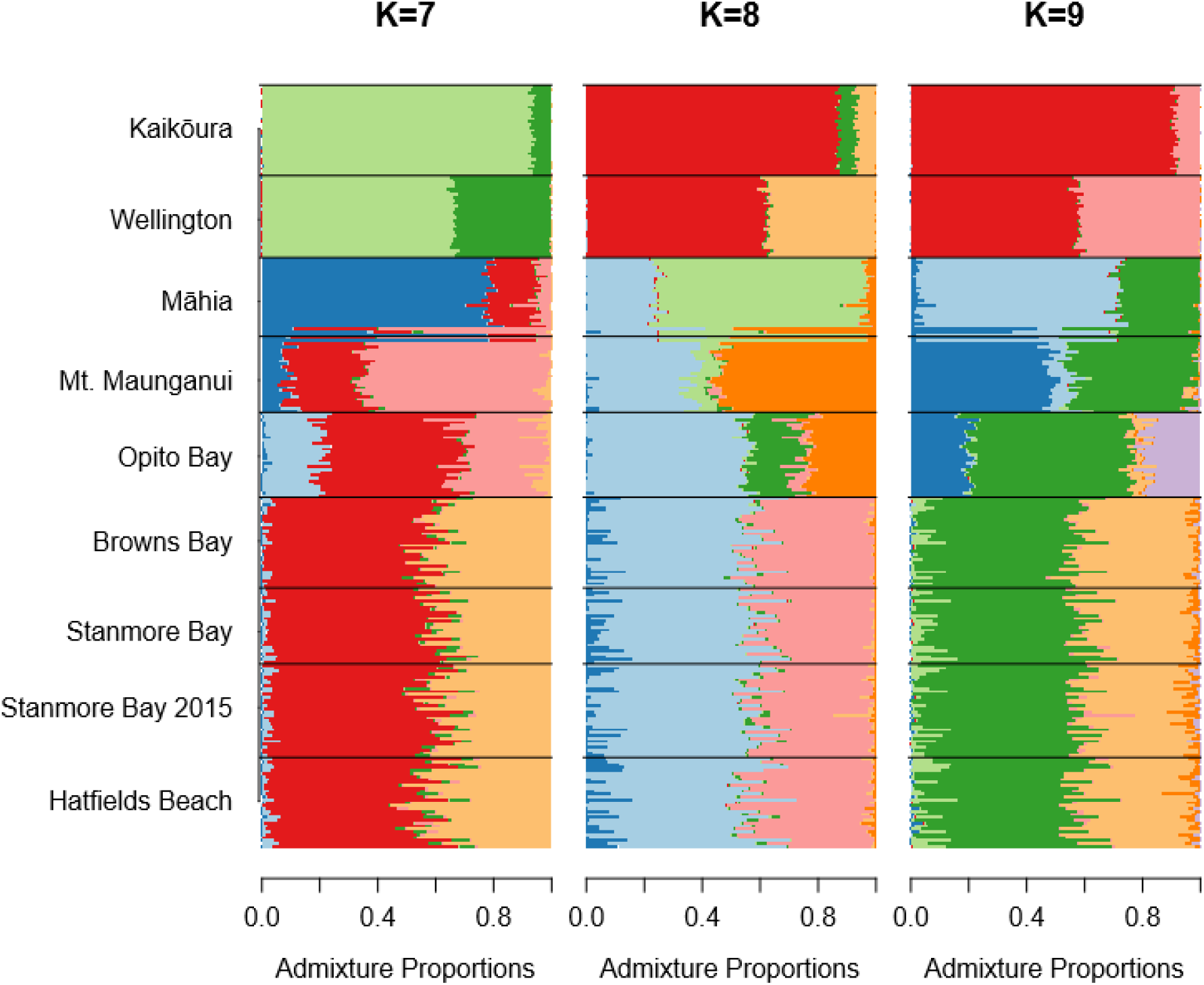
Admixture plots based on Bayesian clustering analyses generated in STRUCTURE for K 7-9. Relatively little structure is observed, and a much higher amount of noise is observed as K increases. In all instances, Kaikoura and Wellington retain a high degree of separation from other peoples, and Māhia also retains some distinctiveness.

